# The effect of common paralytic agents used for fluorescence imaging on redox tone and ATP levels in *Caenorhabditis elegans*

**DOI:** 10.1101/2023.09.21.558750

**Authors:** Katherine S Morton, Ashlyn K Wahl, Joel N Meyer

## Abstract

One aspect of *Caenorhabditis elegans* that makes it a highly valuable model organism is the ease of use of *in vivo* genetic reporters, facilitated by its transparent cuticle and highly tractable genetics. Despite the rapid advancement of these technologies, worms must be paralyzed for most imaging applications, and few investigations have characterized the impacts of common chemical anesthetic methods on the parameters measured, in particular biochemical measurements such as cellular energetics and redox tone. Using two dynamic reporters, QUEEN-2m for relative ATP levels and reduction-oxidation sensitive GFP (roGFP) for redox tone, we assess the impact of commonly used chemical paralytics. We report that no chemical anesthetic is entirely effective at doses required for full paralysis without altering redox tone or ATP levels, though 100 mM 2,3-Butadione monoxime appears to be the least problematic. We also assess the use of cold shock, commonly used in combination with physical restraint methods, and find that cold shock does not alter either ATP levels or redox tone. In addition to informing which paralytics are most appropriate for research in these topics, we highlight the need for tailoring the use of anesthetics to different endpoints and experimental questions. Further, we reinforce the need for developing less disruptive paralytic methods for optimal imaging of dynamic *in vivo* reporters.

## 1. Introduction

*Caenorhabditis elegans* is a powerful model organism widely used by researchers to study developmental biology, neurobiology, metabolism, and human diseases such as cancer, metabolic disorders, and neurodegeneration (1, 2). Worms reach adulthood in just 3 days, are highly genetically tractable, and inexpensive to maintain. Of particular utility for *C. elegans* researchers is their transparency, making them ideal for microscopy-based investigations.

Ranging from assessing localization and morphology, to gene expression via production of gene-specific promoter-driven fluorescent “reporter” proteins, to dynamic real time read-outs of biochemical endpoints such as Ca^2+^ dynamics, redox tone, and ATP levels, light microscopy is at the forefront of methods used in *C. elegans* research (3). The generation of real-time *in vivo* reporters that allow for fluorescent readouts of biochemical endpoints offers large advantages over traditional biochemical assessments, but their utility may be limited if the chemical paralytics typically used for imaging alter the biochemical endpoints being measured. Unlike transcriptional reporter strains, these sensors respond rapidly to intracellular conditions, including changes that may be introduced by anesthetics. Two areas of sensor development that have driven significant advancement aim to assess reactive oxygen species or redox tone and ATP levels.

Reactive oxygen species (ROS) are various products of the reduction of oxygen from the addition of electrons; they serve necessary signaling roles at basal levels, but at high levels they can oxidize DNA, proteins, and lipids resulting in cellular damage (4). In most cells, mitochondria are the primary producers of ROS via the electron transport chain (4). Historically, detecting alterations to reactive oxygen species production in *C. elegans* has been achieved through quantification of GFP driven by promoters of antioxidant genes, quantifying byproducts of oxidation (oxidized lipids, DNA damage), or the use of dyes (5). However, these methods can lack specificity and sensitivity, are difficult to quantify in individual compartments, or are not dynamic enough to demonstrate rapid changes in redox tone. More recently, fluorescent sensors such as Hyper and reduction:oxidation sensitive GFP (roGFP) have allowed for compartment-specific expression and assessment of response in real time (6).

Similarly, cellular regulation of ATP levels can now be dynamically monitored *in vivo* through reporters such as PercevalHR and QUEEN-2m (7, 8). Produced through glycolysis and oxidative phosphorylation, ATP levels and ATP:ADP ratio supply valuable insight into bioenergetic function and can indicate differences in metabolite availability, TCA cycle and glycolytic function, and overall cellular bioenergetic homeostasis. In the absence of *in vivo* reporters, ATP must be measured quantitatively at the whole worm level, or after cell sorting. Dynamic by design, ATP levels change rapidly rendering analyses that require more invasive methods less accurate. The use of fluorescent reporters bypasses this issue at the cost of requiring paralysis or immobilization for effective imaging.

Though critical to accurate reporting and interpretation of these dynamic fluorescent sensors, no document currently exists that comprehensively reports the effects of common anesthetics on *C. elegans* cellular redox tone and ATP levels. Whether mounted on slides or imaged in multiwell plates, most common imaging techniques require the full paralysis of nematodes to be effective. Historically, and most commonly today, this is conducted with chemical paralytic agents. As each anesthetic has a discrete mechanism by which paralysis is induced, we evaluated 5 different chemical paralytic agents as well as cold shock.

Likely the most common paralytic, sodium azide is a well-established inhibitor of mitochondrial respiration that blocks cytochrome c oxidase (electron transport chain Complex IV) by binding to the oxygen reduction site (9, 10). Paralysis from exposure to sodium azide is generally ascribed to the rapid depletion of ATP necessary for movement. Azide also inhibits ATP hydrolysis by F_1_-ATPases, leading to stimulation of potassium channels in bovine *in vitro* models (11). ATP-gated potassium channels are essential for coupling cellular ATP levels to membrane excitability and represent a potential contribution to both paralysis and off-target effects (12). Sodium azide use in *C. elegans* was previously not found to induce the stress response pathways associated with *hif-1, TMEM-135, hsp-4, hsp-16.2*, or *gcs-1* (13). It has been used as a paralytic agent for a wide range of studies, including neurotoxicology, gravitaxis and oxidative stress (14-16).

Levamisole HCl has long been used as an anti-helminthic and is used in *C. elegans* not only as a paralytic, but to examine anthelminthic resistance. Levamisole binds to and activates L-type acetylcholine receptors leading to sustained Ca^2+^ flux into neurons and muscles, ultimately inducing spastic paralysis (9-11).

2,3-Butanedione monoxime (2,3-BDM) is an inhibitor of skeletal muscle myosin II, that specifically inhibits the myosin ATPase activity. This inhibition causes a decrease in muscle force production (12). A recent investigation of the contribution of the reproductive system to oxygen consumption in *C. elegans* utilized 2,3-BDM, stating that this method of inhibition is “not increasing ATP synthesis or mitochondrial oxygen consumption” (13); however, to our knowledge, no studies have empirically tested the effect of 2,3-BDM on ATP levels. 2,3-BDM induces flaccid paralysis, such that muscles are not contracted during imaging, but this effect appears to be relatively slow; previous work often allows the worms a full hour to paralyze prior to imaging (13-15).

The exact mechanism of action of 1-phenoxy-2-propanol (1P2P) remains unknown, but it has been shown to eliminate action potentials in neurons and reduce muscle contraction excitability (24). Despite lack of a clear mechanism, 1P2P exposure induces ATP depletion in *C. elegans*, and both ATP depletion and collapse of mitochondrial membrane potential in Neuro2a cells (25). Despite limited evidence, these data suggest mitochondria as a target for the mechanism of 1P2P. Mammalian toxicology evaluations find no risk of skin irritation and low levels of mucosal irritation suggesting low overt toxicity in mammalian models; however, mitochondrial endpoints were not evaluated (26, 27).

Finally, cold shock is used in combination with physical impediments to movement such as polystyrene beads. Exposure to cold temperature slows metabolic processes in the worm, reducing movement. Cold shock at 2 C was only lethal after 12-hour exposures or longer, though cold shocked worms show decreased gonad size, loss of pigmentation, and increased vulval abnormalities after recovery (16). Cold shock at 4 C for 16 hours is capable of inducing blebbing of dopaminergic neurons, suggesting it may induce dopaminergic neurodegeneration (17). As most paralytic uses of cold shock are far shorter in duration, it is unclear whether these effects occur in this timeframe.

Previous work has reviewed the time required to induce paralysis, recovery from paralysis, and induction of a number of stress-responsive genes via GFP reporter constructs after exposure to-phenoxy-2-propanol, sodium azide, levamisole HCl, and cold shock (18). However, the real-time impacts of these and additional paralytics on more dynamic reporters have not been assessed.

Here, we investigate how quickly the common anesthetics sodium azide, levamisole HCl, 1-phenoxy-2-propanol (1P2P), 2,3-butadione monoxime (2,3-BDM), and 4 C cold shock induce paralysis, how quickly the worms can recover from exposure, and the effects of the treatments on two key biochemical parameters measured by fluorescent reporters: cellular redox tone, in this case assessed as the oxidation status of the glutathione pool of all cells (roGFP) (19) and relative ATP levels in muscle cells (QUEEN-2m)(20). We report that all chemical paralytics, though not acute cold shock, induce changes to either redox tone or ATP levels at high doses typically used for effective paralysis and imaging, and highlight the need for tailoring anesthetic use to the experimental questions being used, or developing more viable non-chemical alternatives.

## 2. Methods

### 2.1 C. elegans Strains and Culturing

*C. elegans* were grown and maintained at 20 C on 10 cm K-agar plates seeded with OP50 *E. coli (21)*. To synchronize worms’ growth and development for experiments, adult worms were transferred to a fresh plate and allowed to lay eggs for two hours. After this time, all worms were rinsed from the plate until only the eggs remained. Eggs were allowed to mature for 72 hours before use in experiments (day 1 of adulthood). Strains used in this study were JV2, GA2001, and N2, all of which were obtained from the *C. elegans* Genome Center.

### 2.2. Drugs and Concentrations

All paralytics were prepared in K-medium (21) at the following concentrations: 10mM, 100mM, and 500mM sodium azide, 100mM and 300mM 2,3-butanedione monoxime, 1mM and 3mM levamisole HCl, and 0.5% and 1.0% 1-phenoxy-2-propanol. The cold shock temperature was 4ºC. Concentrations were selected based on commonly used doses in cited *C. elegans* literature. If only one dose is commonly used, such as 1% for 1P2P, a half dose was examined to provide a wider scope of evaluation for levels without off target effects.

### 2.3. Assessment of Paralysis

To assess paralysis, ten 20 μL drops of the given drug and concentration were placed in a small petri dish. One N2 worm was transferred into each drop by flame sterilized platinum pick. The time it was placed was noted, and the worm was checked every 60 seconds for the first five minutes, and then every five minutes until the worm was paralyzed, or until thirty minutes had passed. Paralysis was defined as no movement when the worm was touched with a pick. The time between when the worm was first placed in the drug droplet and when it reached paralysis was recorded. Time to 50% paralysis was determined by performing a Cumulative Gaussian non-linear fit in GraphPad Prism 9.3.0. The mean and standard deviation provided are best-fit values.

### 2.4. Assessment of Recovery from Paralysis

To quantify recovery of the worms from paralysis by each agent, twenty worms were transferred into a 1.5 mL Eppendorf tube containing 400 μL of the given drug and concentration. After a 30-minute exposure, the drug was aspirated from the tube, and the worms were rinsed with K-medium three times. The worms were centrifuged for thirty seconds at 2200 rpm between each rinse to minimize loss of worms. In the event the worms stuck to the sides of the tube, 10 μL of 0.1% triton were added to the solution before rinsing.

After washing, exposed worms were placed onto a 10 cm K-agar plate seeded with OP50 *E. coli*. The worms were then observed every fifteen minutes for two hours. With each observation, the number of worms that moved when touched with a pick, or were moving on their own, was recorded.

### 2.5. Assessment of cytosolic redox tone and ATP levels

Synchronized adult JV2 or GA2001 worms were assessed using a FLUOstar Omega microplate reader (BMG Labtech) with each well containing 1000 worms suspended in 100 uL of K-medium. Just before insertion into the plate reader, 100 μL of 2X anesthetic or H_2_O_2_ was added to the wells (final concentration of 1X). Fluorescence was recorded every 5 minutes for 35 minutes at 405 nm and 485 nm excitation wavelengths with a 520 nm emission wavelength.

To analyze the data, the average fluorescence of blank wells corresponding to each treatment was subtracted from the fluorescence of each corresponding well containing worms at each wavelength, which could then be converted to ratios of one excitation wavelength to another. This process was completed for each time point and each treatment group. For JV2 (roGFP) ratios are reported as oxidized: reduced, representing 485 nm excitation:405 nm excitation. For GA2001 (Queen-2m) ratios are reported as 405 nm:485 nm, representing ATP-bound:unbound protein.

### 2.6. Statistical Analysis

Statistical analysis for all data was completed with GraphPad Prism 9.3.0. All data were assessed by a Shapiro-Wilks Normality test, followed by a 2-way Kruskal-Wallis as no data were normally distributed. Post-hoc tests consisted of Dunnett’s multiple comparisons tests. P<0.05 was considered statistically significant, and all error bars reflected the standard error of the mean. Key statistical comparisons are described in the Results section, and full statistical analysis p-values are listed in Supplemental Tables 1-3.

## 3. Results

### Sodium azide rapid paralyzes while decreasing both ATP levels and redox tone

In this study we examined sodium azide at three concentrations: 10 mM, 100 mM, and 500 mM, and report that all sodium azide concentrations induce 50% population paralysis in less than a minute, ultimately resulting in paralysis of the entire population (Fig 1A). Though 10 mM and 100 mM exposures are 76.6% and 85.0% recoverable within 2 hours, this decreases dramatically to 1.8% at 500 mM (Fig 1A). All concentrations of sodium azide result in a rapid depletion of ATP in muscle cells that remains stable for at least 30 mins (Fig 1B), consistent with the role of azide as an inhibitor of oxidative phosphorylation (22, 23). Similarly, all concentrations of sodium azide cause a temporary decrease in cytosolic redox tone (i.e., more reduced state) of the glutathione pool (Fig 1C).

**Fig 1.**
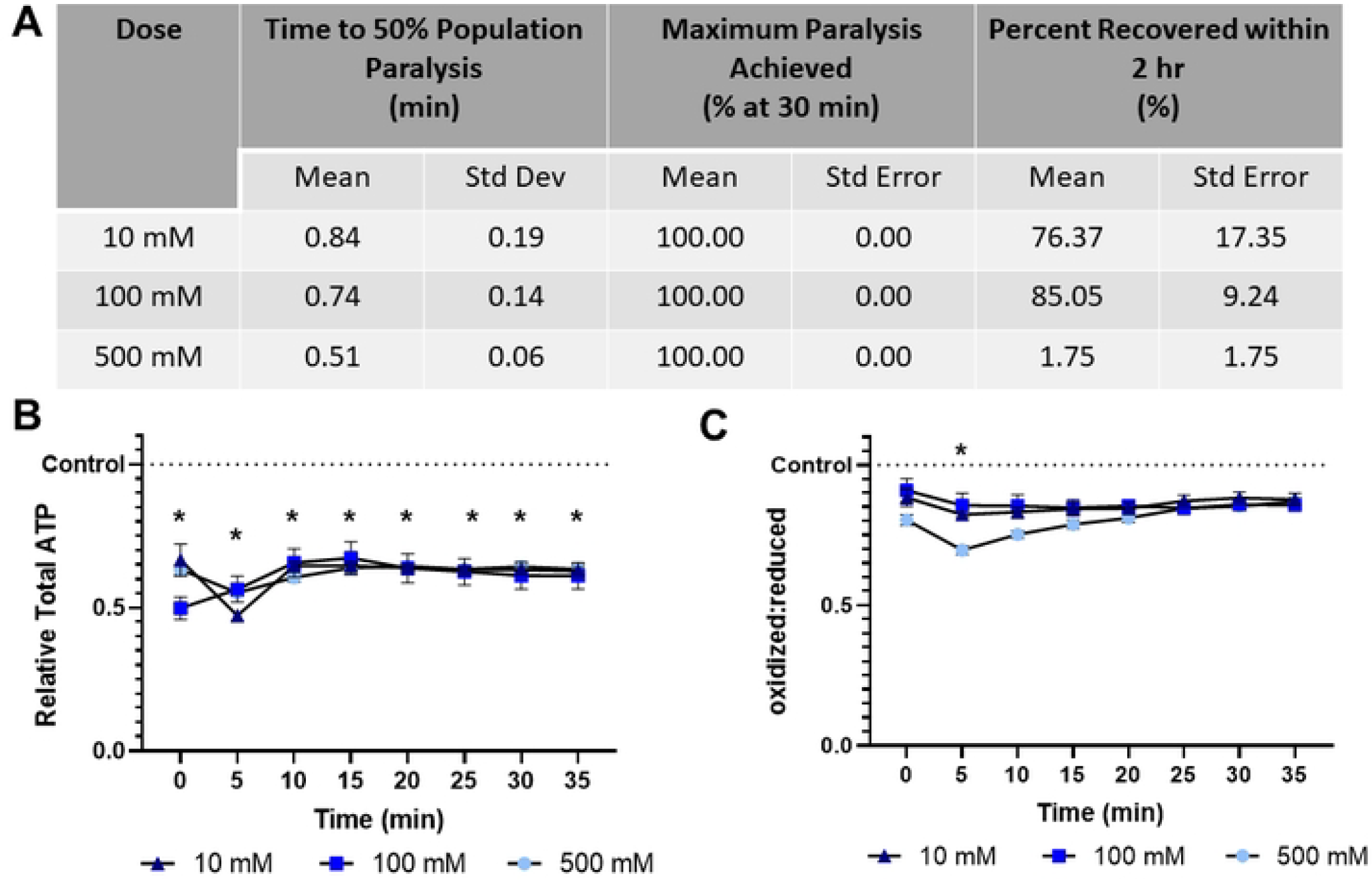
Sodium azide induces rapid paralysis, ATP depletion, and briefly decreases glutathione pool oxidation **(A)** The time required for 10 mM, 100 mM, and 500 mM sodium azide to induce paralysis, the maximum percentage of worms paralyzed and the percentage that recover within 2 hours was determined. **(B)** Relative total muscle cell ATP levels were quantified for 35 minutes using the fluorescent reporter p*myo-3*::Queen-2m after exposure to 10 mM, 100 mM, and 500 mM sodium azide respectively. Statistical analysis was performed by two-way Kruskal-Wallis, with Dunnett’s post-hoc. **(C)** Redox tone of the cytosolic glutathione pool was assessed through a reduction:oxidation sensitive GFP every 5 minutes for 35 minutes after exposure to 10 mM, 100 mM, and 500 mM sodium azide respectively. Statistical analysis was performed by two-way Kruskal-Wallis, with Dunnett’s post-hoc.

### Levamisole HCl induces paralysis and ATP depletion without alteration to cellular redox tone

Levamisole HCl is slower to paralyze than sodium azide, requiring 15.69 and 9.26 minutes to achieve 50% population paralysis at 1 mM and 3 mM respectively (Fig. 2A). Further, only 70% and 80% of the population was paralyzed within 30 mins. No individuals recovered within 2 hours, in agreement with previous reports stating that over 4 hours are required for recovery (18). Despite inducing ATP depletion, unlike azide, levamisole did not modify cytosolic redox tone (Fig. 2A, 2B).

**Fig 2.**
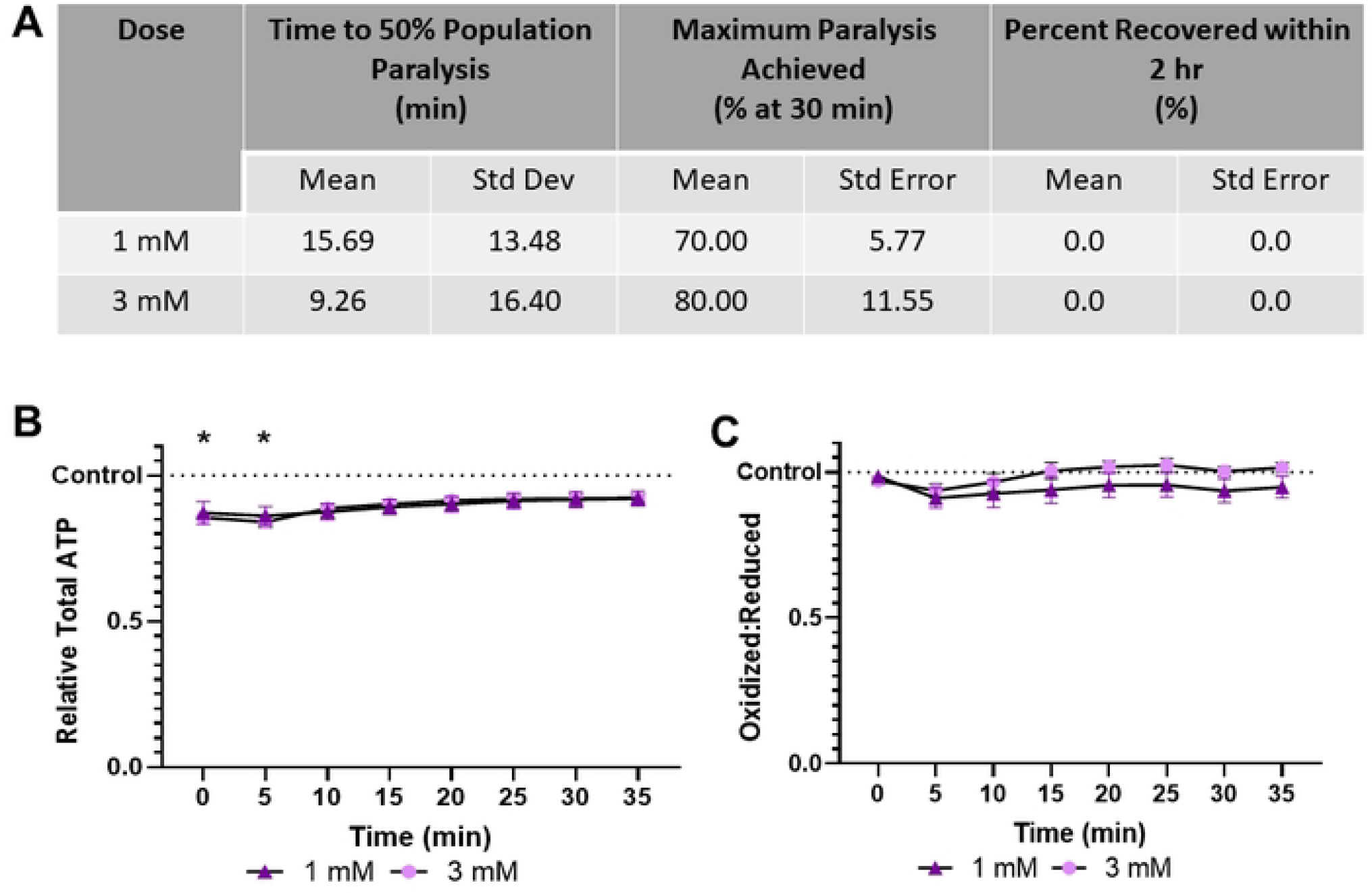
Levamisole HCl induces paralysis and mild ATP depletion, without altering glutathione pool oxidation **(A)** The time required for 1 mM and 3 mM Levamisole HCl to induce paralysis, the maximum percentage of worms paralyzed and the percentage that recover within 2 hours was determined. **(B)** Relative total muscle cell ATP levels were quantified for 35 minutes using the fluorescent reporter p*myo-3*::Queen-2m after exposure to 1 mM and 3 mM Levamisole HCl respectively. Statistical analysis was performed by two-way Kruskal-Wallis, with Dunnett’s post-hoc. **(C)** Redox tone of the cytosolic glutathione pool was assessed through a reduction:oxidation sensitive GFP every 5 minutes for 35 minutes after exposure to 1 mM and 3 mM Levamisole HCl respectively. Statistical analysis was performed by two-way Kruskal-Wallis, with Dunnett’s post-hoc.

### 1P2P exposure causes high oxidative stress and ATP depletion

1P2P, at 0.5% and 1.0%, rapidly paralyzes worms in less than 3 minutes and achieves 100% paralysis (Fig. 3A). However, it induces ATP depletion that progressively increases over time, and high elevation of the cytosolic redox tone, indicating an increasingly oxidative cellular environment (Fig. 2A, 2B). Notably, 100% and 35.85% of the nematodes tested recover within 2 hours at the high and low dose, demonstrating the ATP depletion and redox stress observed are not immediately lethal.

**Fig 3.**
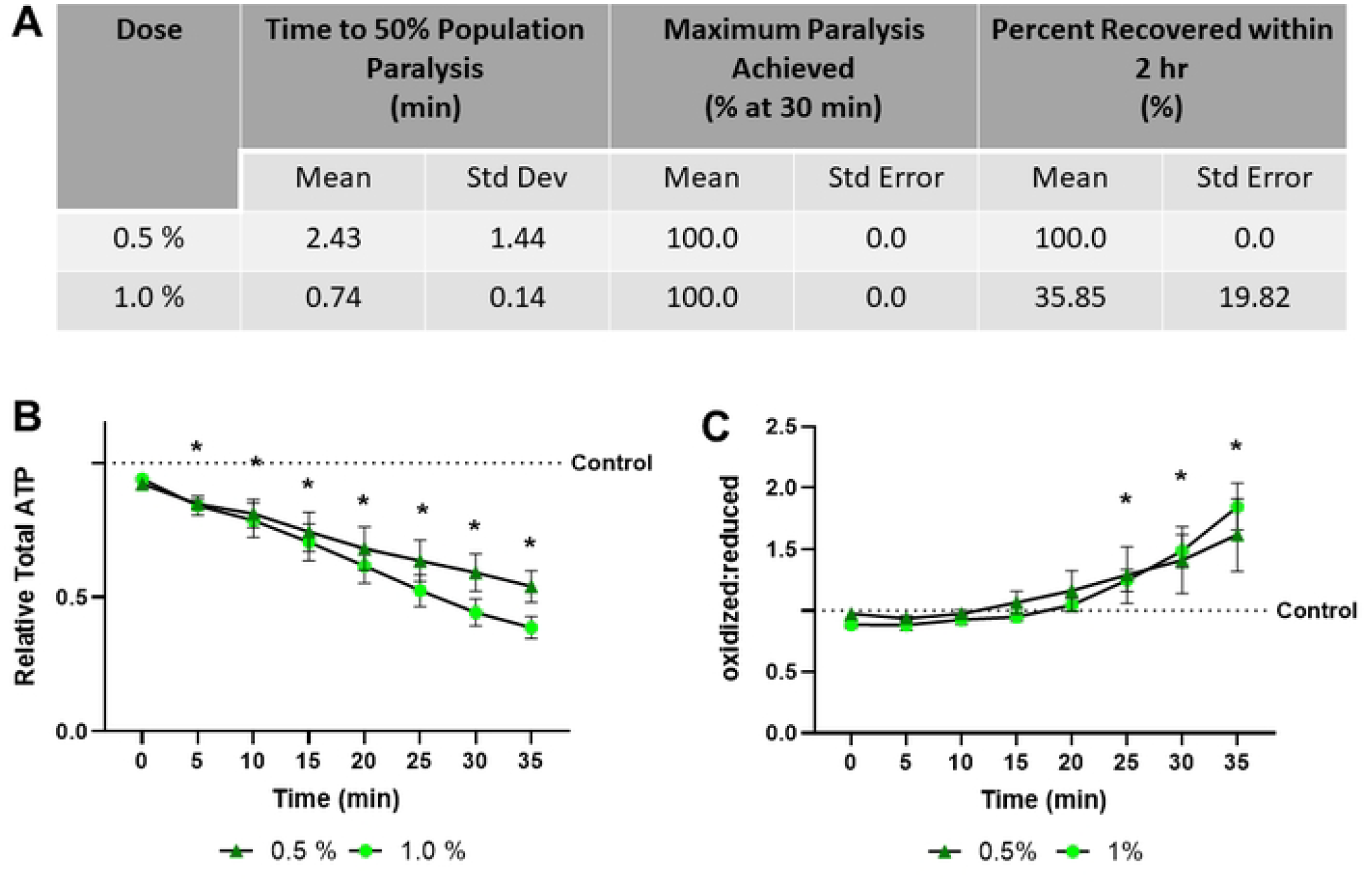
1-Phenoxy-2-propanol exposure results in rapid paralysis, severe ATP depletion, and swift oxidation of the glutathione pool **(A)** The time required for 0.5% and 1.0% 1-phenoxy-2-propanol to induce paralysis, the maximum percentage of worms paralyzed and the percentage that recover within 2 hours was determined. **(B)** Relative total muscle cell ATP levels were quantified for 35 minutes using the fluorescent reporter p*myo-3*::Queen-2m after exposure to 0.5% and 1.0% 1-phenoxy-2-propanol respectively. Statistical analysis was performed by two-way Kruskal-Wallis, with Dunnett’s post-hoc. **(C)** Redox tone of the cytosolic glutathione pool was assessed through a reduction:oxidation sensitive GFP every 5 minutes for 35 minutes after exposure to 0.5% and 1.0% 1-phenoxy-2-propanol respectively. Statistical analysis was performed by two-way Kruskal-Wallis, with Dunnett’s post-hoc.

### 2,3-BDM is effective without clear toxicity at 100 mM, but lethal at 300 mM

2,3-BDM shows divergence between the examined concentrations. 100 mM results in slow paralysis of 83% of the population, but high recovery (85.84%) and no significant effect on ATP levels or redox tone (Fig. 4A). Conversely, 300 mM 2,3-BDM rapidly depletes ATP levels and increases redox tone as it paralyzes 50% of the population in less than 3 minutes. The increase in redox tone appears to be as severe as treatment with 3% hydrogen peroxide (Fig. 4B), a dose lethal within 1 hour. 300 mM 2,3-BDM was capable of inducing the “blue wave of death” (24) in previous experiments, supporting the notion that it is lethal within 30 mins (data not shown).

**Fig 4.**
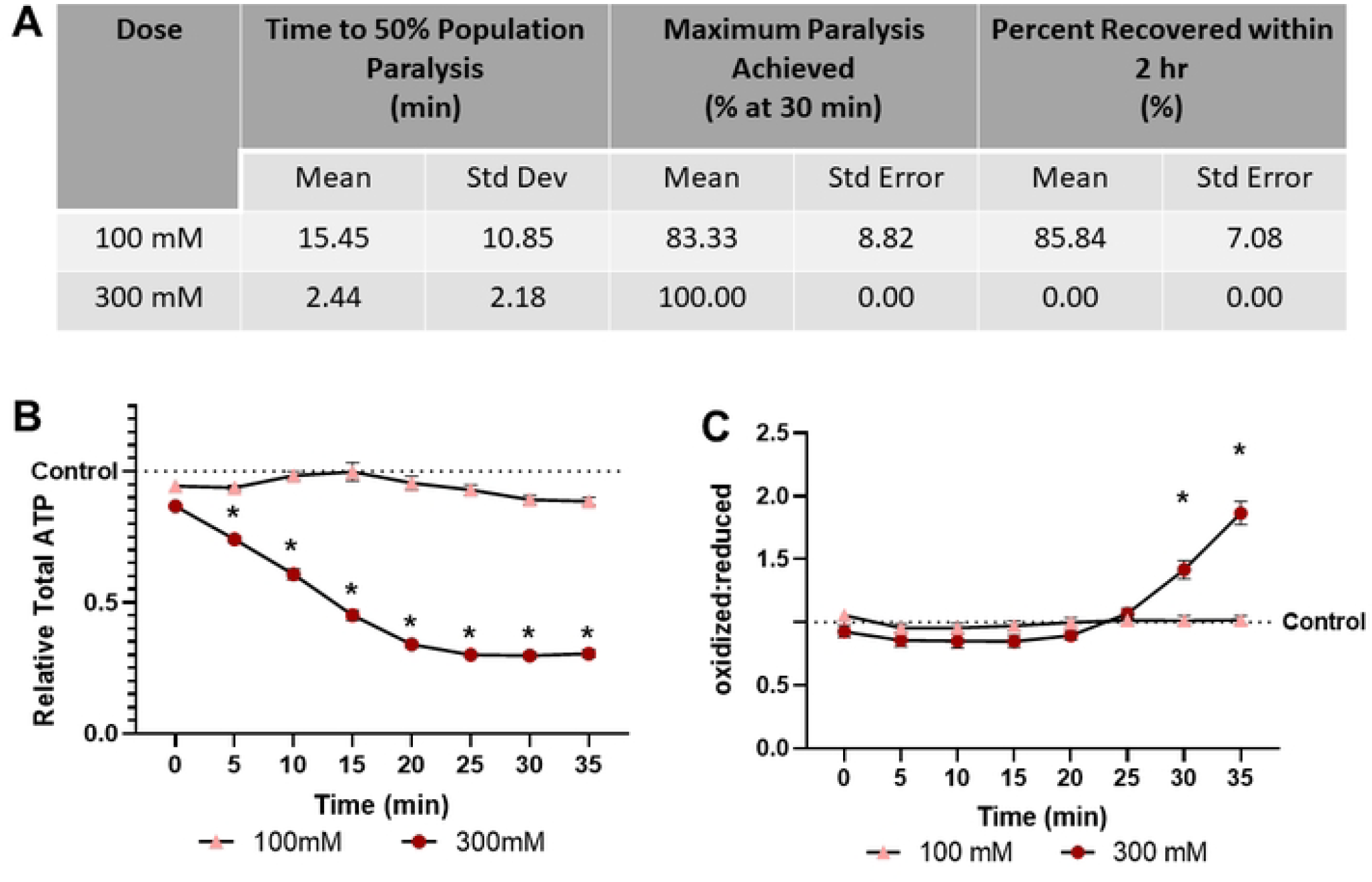
2,3-Butadione monoxime treatment results in dose dependent paralysis, ATP depletion, and oxidation of the glutathione pool **(A)** The time required for 100 mM and 300 mM 2,3-butadione monoxime to induce paralysis, the maximum percentage of worms paralyzed and the percentage that recover within 2 hours was determined. **(B)** Relative total muscle cell ATP levels were quantified for 35 minutes using the fluorescent reporter p*myo-3*::Queen-2m after exposure to 100 mM and 300 mM 2,3-butadione monoxime respectively. Statistical analysis was performed by two-way Kruskal-Wallis, with Dunnett’s post-hoc. **(C)** Redox tone of the cytosolic glutathione pool was assessed through a reduction:oxidation sensitive GFP every 5 minutes for 35 minutes after exposure to 100 mM and 300 mM 2,3-butadione monoxime respectively. Statistical analysis was performed by two-way Kruskal-Wallis, with Dunnett’s post-hoc.

### Acute cold shock does not paralyze, alter ATP levels, or change redox tone in worms

Finally, 30-minute 4°C cold shock alone is insufficient to paralyze more than 15% of the population (Fig. 5A). This is anticipated as cold shock is typically utilized in combination with physical barriers or microfluidics. Importantly, cold shock did not detectably alter either muscular ATP levels or cytosolic glutathione pool redox tone, supporting its use over chemical paralytics from the perspective of disturbing these parameters (Fig. 5A, 5B).

**Fig 5.**
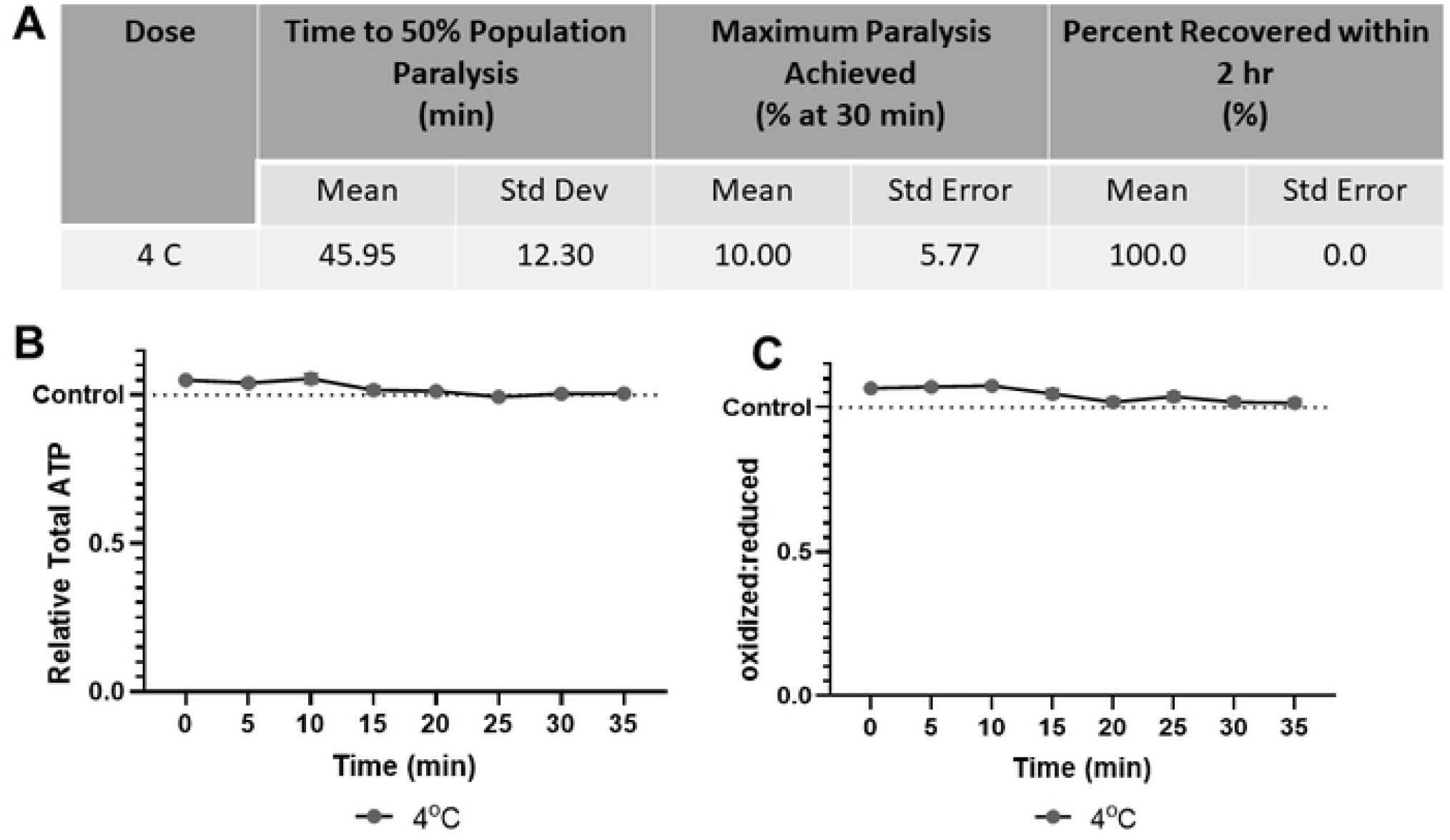
4° C Cold shock does not paralyze worms, alter muscular ATP level, or glutathione pool redox tone **(A)** The time required for 4° C cold shock to induce paralysis, the maximum percentage of worms paralyzed and the percentage that recover within 2 hours was determined. **(B)** Relative total muscle cell ATP levels were quantified for 35 minutes using the fluorescent reporter p*myo-3*::Queen-2m after exposure to 4° C cold shock. Statistical analysis was performed by two-way Kruskal-Wallis, with Dunnett’s post-hoc. **(C)** Redox tone of the cytosolic glutathione pool was assessed through a reduction:oxidation sensitive GFP every 5 minutes for 35 minutes after exposure to 4° C cold shock. Statistical analysis was performed by two-way Kruskal-Wallis, with Dunnett’s post-hoc.

### Combined comparison of all methods

To facilitate direct comparison of all paralytic methods tested, we present graphs combining all time-to-paralysis results (Fig. 6A), time to recovery results (Fig. 6B), effects on ATP (Fig. 6C), and effects on redox state (Fig. 6D).

**Fig 6.**
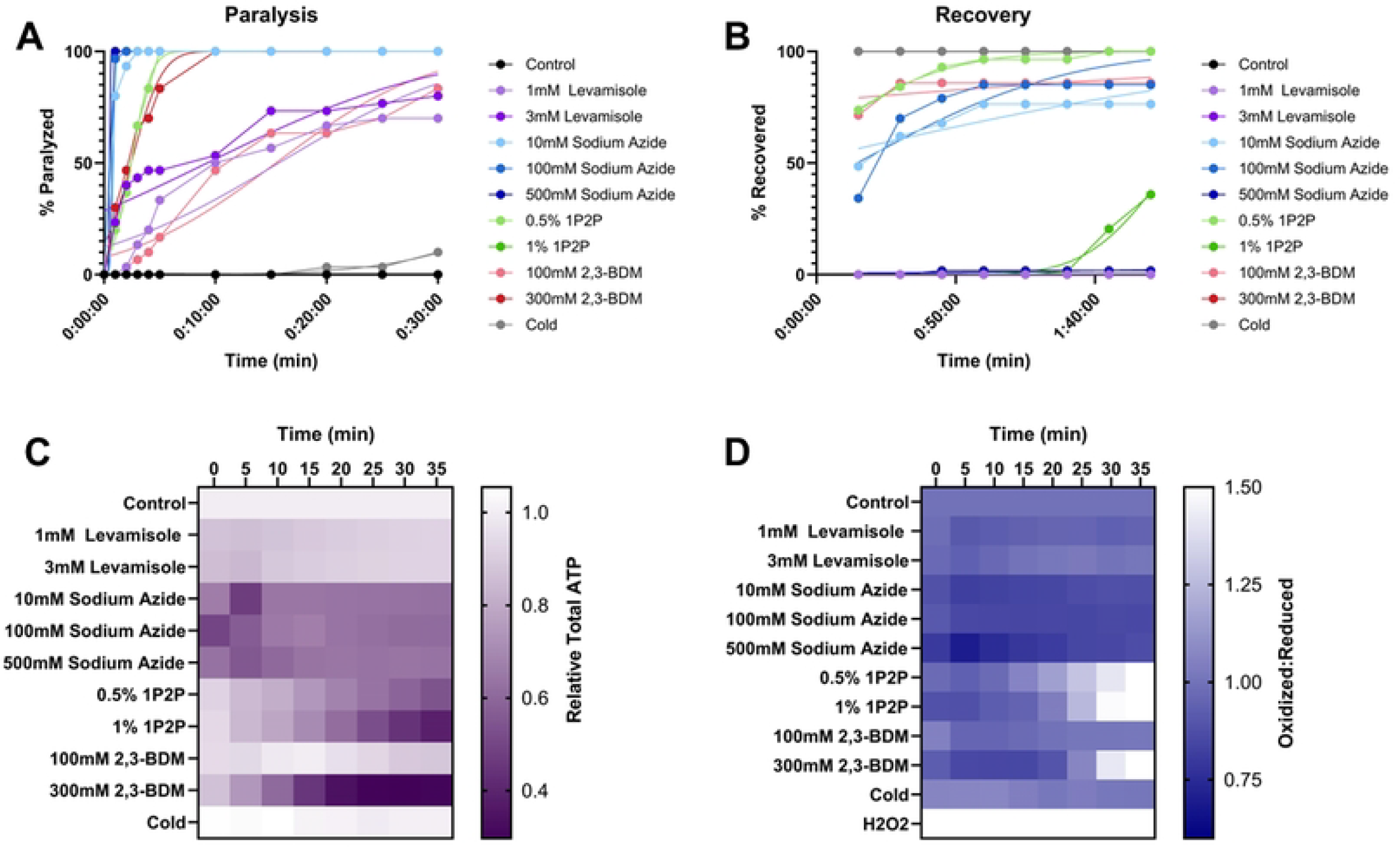
No common chemical *C*. *elegans* anesthetic results in greater than 50% population paralysis without alteration to muscular ATP levels or cytosolic redox tone **(A)** The percentage of worms paralyzed over time after exposure to an anesthetic. **(B)** The percentage of worms recovered from anesthetic exposure across a 2-hour monitoring window. **(C)** The evaluation of common anesthetics and doses to alter intra-muscular ATP levels was quantified every 5 minutes for 35 minutes with a p*myo-3*::Queen-2m fluorescent reporter. All anesthetics and doses were compared to non-treated controls at each time point through a 2-way Kruskal-Wallis with Dunnett’s post hoc. **(D)** Redox tone of the cytosolic glutathione pool was assessed through a reduction:oxidation sensitive GFP every 5 minutes for 35 minutes after injection of common anesthetics. Statistical analysis was performed by two-way Kruskal-Wallis, with Dunnett’s post-hoc. Each dose of each anesthetic was compared to controls at each timepoint.

## 4. Discussion

In this work, we report the impact of 4 chemical anesthetics and a 4° C cold shock on intramuscular cytosolic ATP levels and cytosolic redox tone of the glutathione pool (in all cells) using the dynamic *in vivo* reporters roGFP and QUEEN-2m in *C. elegans*. Due to the rapid equilibrium between mitochondria and the cytosol for both adenylate nucleotides and glutathione oxidation, these cytosolic reporters allow key insight into the impact of common anesthetics on mitochondrial function (25, 26). Investigating how long it took for worms to paralyze for each given treatment revealed that all treatments studied, besides cold shock, lead to statistically significant paralysis in five minutes or less. 10 mM, 100 mM, and 500 mM sodium azide and 1% 1-phenoxy-2-propanol resulted in paralysis the fastest; however, sodium azide resulted in less ATP depletion and much less redox stress, making it the more appealing paralytic for rapid paralysis. However, as sodium azide still decreased redox tone and decreased ATP levels, it remains an imperfect paralytic for mitochondrial or redox related endpoints. The only chemical paralytic agent that does not decrease ATP or alter cytosolic redox tone is levels is 100 mM 2,3-BDM. However, it requires more than 15 minutes to achieve 50% population paralysis, decreasing its utility for measurements that need to be taken rapidly. Finally, levamisole HCl is the only paralytic that never induces alterations to redox tone, supporting its use for redox related endpoints.

All of the drugs and paradigms tested here or in previous work resulted in at least 10% recoverable worms within 2 hours of exposure, with the exceptions of 500 mM sodium azide and 300 mM 2,3 BDM. The lack of a fluorescent wave in sodium azide imaging (data not shown) suggests it is not lethal, unlike 300 mM 2,3-BDM. Prior evaluations utilizing doses between 295-300 mM 2,3-BDM report requiring 30 minutes to a full hour before assessing their respective endpoint (13, 15, 27), in which case the worms may be deceased upon imaging. Especially given the lack of mechanistic explanation for the lethality of 300 mM 2,3-BDM, dose responses should be performed to determine acceptable dose ranges for future experiments. The lethality of an anesthetic that has been widely used highlights the need for pursuit of more ideal anesthetics or alternative methods.

The desire to pursue non-chemical anesthetics has led to a boom in the development of microfluidic devices, physical barriers such as polystyrene beads, and polymers that can be hardened after the addition of worms, such as BIO-133. As these methods rely on friction that the worms cannot overcome to move (28), loading the worms into narrow channels (29), or manipulation with acoustic waves (30, 31), we predict that they are less likely to alter bioenergetics and redox tone. The ultraviolet (365 nm) light curable polymer BIO-133 (32, 33) would be expected to induce DNA damage, though to our knowledge this has not been investigated. The use of methods that do not require hands-on dosing have also been employed in “lab-on-a chip” models to produce high-throughput readouts, pushing forward the ability to use *C. elegans* for chemical screening without anesthetic interference (34).

The lethality of 300 mM 2,3-BDM, as well as the impacts of other anesthetics on roGFP and Queen-2m readout, demonstrate the critical need to tailor anesthetic use to experimental methods and goals. Beyond our evaluation, the impacts of the examined anesthetics may vary based on variation in exposure method, size of exposure groups (worms per amount of toxicant), and exposure conditions (liquid vs solid media). Additionally, uptake of the drugs likely varies depending on life stage as the protective cuticle of the worm thickens with age, altering the amount of drug that can penetrate (35). This was not an exhaustive study of all possible mitochondrial parameters, and the potential that the mitochondria are being impaired by the chemical in ways that the roGFP and Queen-2m assays are not sensitive to should be considered. However, our findings lay the groundwork for understanding which anesthetics are acceptable for bioenergetic and redox related endpoints.

## 5. Acknowledgements

Some strains were provided by the *Caenorhabditis* Genetics Center, which is funded by NIH Office of Research Infrastructure Programs (P40 OD010440). This work was supported by National Institute of Health (P42ES010356, R01ES034270, 2R44OD024963, and T32ES021432).

## Notes

### Competing Interest Statement

The authors have declared no competing interest.

